## Supplementary Information for "The energy landscape reshaped by strain-specific mutations underlies the long-range epistasis in NS1 evolution of influenza A virus"

**Table S1.** X-ray diffraction and refinement statistics of the complexes of VN NS$^{ED}$ and p85β$^{iSH2}$.

| | **VN NS1-ED W187A:p85β complex** PDB ID: 7RCH |
|---|---|
| **Data collection*** | |
| Space group | $P2_1$ |
| Cell dimensions | |
| *a*, *b*, *c* (Å) | 61.44, 92.84, 67.12 |
| α, β, γ (°) | 90.00, 105.03, 90.00 |
| $R_{merge}$ | 0.169 (0.720) |
| $R_{pim}$ | 0.106 (0.442) |
| $I / \sigma I$ | 4.0 (1.3) |
| Completeness (%) | 97.8 (96.8) |
| Redundancy | 3.5 (3.6) |
| **Refinement** | |
| Resolution (Å) | 3.1 (3.31 to 3.10)** |
| No. Reflections | 13026 |
| $R_{work}/R_{free}$ | 0.243/0.276 |
| No. atoms | |
| Protein | 4204 |
| Water | 0 |
| *B*-factors | |
| Overall | 74.29 |
| R.m.s. deviations | |
| Bond lengths (Å) | 0.002 |
| Bond angles (°) | 0.447 |

* The numbers in parentheses refer to the highest resolution shell.

** A single crystal was used to collect each of the dataset.

Table S2. Chemical shifts of 1918 NS1 methyl atoms.

| Residues | Atom name | $^{13}C$ | $^{1}H$ | Residues | Atom name | $^{13}C$ | $^{1}H$ |
|---|---|---|---|---|---|---|---|
| A79 | CB-HB | 19.52 | 1.530 | L144 | CD1-HD1 | 23.68 | 0.774 |
| T80 | CG2-HG2 | 22.00 | 1.181 | L144 | CD2-HD2 | 26.26 | 0.562 |
| I81 | CD1-HD1 | 13.77 | 0.885 | I145 | CD1-HD1 | 12.41 | 0.822 |
| I81 | CG2-HG2 | 17.43 | 0.974 | I145 | CG2-HG2 | 17.52 | 0.925 |
| A82 | CB-HB | 19.53 | 1.372 | L146 | CD1-HD1 | 22.39 | 1.199 |
| V84 | CG1-HG1 | 20.85 | 0.993 | L146 | CD2-HD2 | 26.14 | 1.001 |
| V84 | CG2-HG2 | 20.31 | 0.931 | L147 | CD1-HD1 | 28.94 | 0.705 |
| A86 | CB-HB | 19.50 | 1.312 | L147 | CD2-HD2 | 24.01 | 0.543 |
| L90 | CD1-HD1 | 25.65 | 0.566 | A149 | CB-HB | 20.39 | 0.765 |
| L90 | CD2-HD2 | 23.98 | 0.543 | T151 | CG2-HG2 | 22.97 | 1.240 |
| T91 | CG2-HG2 | 20.74 | 0.874 | A155 | CB-HB | 19.31 | 1.374 |
| M93 | CE-HE | 18.07 | 1.945 | I156 | CD1-HD1 | 15.40 | 0.692 |
| T94 | CG2-HG2 | 21.93 | 1.258 | I156 | CG2-HG2 | 16.63 | -0.753 |
| L95 | CD1-HD1 | 24.12 | 0.904 | V157 | CG1-HG1 | 22.70 | 0.829 |
| L95 | CD2-HD2 | 24.07 | 0.874 | V157 | CG2-HG2 | 18.60 | 0.476 |
| M98 | CE-HE | 17.00 | 1.995 | I160 | CD1-HD1 | 14.20 | 0.708 |
| M104 | CE-HE | 17.65 | 1.535 | I160 | CG2-HG2 | 18.66 | 0.635 |
| L105 | CD1-HD1 | 24.07 | 0.920 | L163 | CD1-HD1 | 26.17 | 0.821 |
| L105 | CD2-HD2 | 24.37 | 0.955 | L163 | CD2-HD2 | N.A. | N.A. |
| M106 | CE-HE | N.A. | N.A. | L166 | CD1-HD1 | N.A. | N.A. |
| V111 | CG1-HG1 | 21.40 | 0.904 | L166 | CD2-HD2 | N.A. | N.A. |
| V111 | CG2-HG2 | 21.52 | 0.816 | T170 | CG2-HG2 | 21.50 | 1.310 |
| A112 | CB-HB | 20.12 | 1.187 | V174 | CG1-HG1 | 24.46 | 0.929 |
| L115 | CD1-HD1 | 26.82 | 0.489 | V174 | CG2-HG2 | 21.09 | 0.624 |
| L115 | CD2-HD2 | 22.87 | 0.273 | A177 | CB-HB | 19.42 | 1.412 |
| I117 | CD1-HD1 | 16.43 | 0.840 | V178 | CG1-HG1 | 24.03 | 0.733 |
| I117 | CG2-HG2 | 19.97 | 0.772 | V178 | CG2-HG2 | 21.79 | 0.467 |
| M119 | CE-HE | 18.07 | 1.815 | V180 | CG1-HG1 | 22.43 | 1.005 |
| A122 | CB-HB | 20.42 | 1.418 | V180 | CG2-HG2 | 21.43 | 0.813 |
| I123 | CD1-HD1 | 9.19 | 0.757 | L181 | CD1-HD1 | 23.48 | 0.798 |
| I123 | CG2-HG2 | 16.66 | 0.804 | L181 | CD2-HD2 | 27.32 | 0.619 |
| M124 | CE-HE | 17.12 | 2.044 | I182 | CD1-HD1 | 13.51 | 0.338 |
| I128 | CD1-HD1 | 14.51 | 0.547 | I182 | CG2-HG2 | 16.79 | 0.726 |
| I128 | CG2-HG2 | 18.68 | 0.631 | L185 | CD1-HD1 | 22.09 | 0.725 |
| I129 | CD1-HD1 | 14.21 | 0.740 | L185 | CD2-HD2 | 26.45 | 0.547 |
| I129 | CG2-HG2 | 17.49 | 0.910 | A187 | CB-HB | 17.89 | 1.523 |
| L130 | CD1-HD1 | 25.44 | 0.659 | T191 | CG2-HG2 | 21.27 | 1.160 |
| L130 | CD2-HD2 | 24.21 | 0.559 | V192 | CG1-HG1 | 22.69 | 0.708 |
| A132 | CB-HB | 25.47 | 1.383 | V192 | CG2-HG2 | 22.82 | 0.670 |
| V136 | CG1-HG1 | 20.55 | 0.398 | V194 | CG1-HG1 | 21.99 | 0.827 |
| V136 | CG2-HG2 | 21.48 | 0.376 | V194 | CG2-HG2 | 20.01 | 0.791 |
| I137 | CD1-HD1 | 13.64 | 0.695 | T197 | CG2-HG2 | 23.83 | 1.389 |
| I137 | CG2-HG2 | 16.81 | 0.744 | L198 | CD1-HD1 | 24.85 | 0.610 |
| L141 | CD1-HD1 | 25.27 | 0.659 | L198 | CD2-HD2 | 23.93 | 0.463 |
| L141 | CD2-HD2 | 26.56 | 0.317 | A202 | CB-HB | 20.30 | 1.017 |
| T143 | CG2-HG2 | 22.35 | 1.097 | | | | |

Table S3. Chemical shifts of PR8 NS1 methyl atoms.

| Residues | Atom name | $^{13}C$ | $^{1}H$ | Residues | Atom name | $^{13}C$ | $^{1}H$ |
|---|---|---|---|---|---|---|---|
| T80 | CG2-HG2 | 21.38 | 1.253 | L144 | CD2-HD2 | 26.16 | 0.557 |
| M81 | CE-HE | 16.81 | 2.083 | I145 | CD1-HD1 | 12.25 | 0.814 |
| A82 | CB-HB | 19.39 | 1.383 | I145 | CG2-HG2 | 17.60 | 0.935 |
| V84 | CG1-HG1 | 20.93 | 0.958 | L146 | CD1-HD1 | 25.84 | 1.025 |
| V84 | CG2-HG2 | 20.35 | 0.927 | L146 | CD2-HD2 | 22.70 | 1.166 |
| A86 | CB-HB | 19.41 | 1.306 | L147 | CD1-HD1 | 28.90 | 0.696 |
| L90 | CD1-HD1 | 25.54 | 0.538 | L147 | CD2-HD2 | 23.90 | 0.540 |
| L90 | CD2-HD2 | 23.85 | 0.502 | A149 | CB-HB | 20.44 | 0.760 |
| T91 | CG2-HG2 | 20.63 | 0.782 | T151 | CG2-HG2 | 22.95 | 1.243 |
| M93 | CE-HE | 17.94 | 1.957 | A155 | CB-HB | 19.28 | 1.373 |
| T94 | CG2-HG2 | 21.85 | 1.249 | I156 | CD1-HD1 | 15.40 | 0.703 |
| L95 | CD1-HD1 | 24.17 | 0.901 | I156 | CG2-HG2 | 16.39 | -0.799 |
| L95 | CD2-HD2 | 24.16 | 0.882 | V157 | CG1-HG1 | 22.67 | 0.829 |
| M98 | CE-HE | 16.97 | 2.002 | V157 | CG2-HG2 | 18.50 | 0.470 |
| M104 | CE-HE | 17.49 | 1.574 | I160 | CD1-HD1 | 14.29 | 0.702 |
| L105 | CD1-HD1 | 24.06 | 0.900 | I160 | CG2-HG2 | 18.30 | 0.657 |
| L105 | CD2-HD2 | 24.02 | 0.854 | L163 | CD1-HD1 | 25.82 | 0.923 |
| I106 | CD1-HD1 | 13.16 | 0.766 | L163 | CD2-HD2 | 23.64 | 0.919 |
| I106 | CG2-HG2 | 17.79 | 0.823 | L166 | CD1-HD1 | N.A. | N.A. |
| V111 | CG1-HG1 | 22.15 | 0.997 | L166 | CD2-HD2 | N.A. | N.A. |
| V111 | CG2-HG2 | 22.04 | 0.961 | T170 | CG2-HG2 | 21.31 | 1.292 |
| A112 | CB-HB | 19.35 | 1.125 | A171 | CB-HB | 18.22 | 1.653 |
| L115 | CD1-HD1 | 26.84 | 0.524 | V174 | CG1-HG1 | 24.18 | 0.909 |
| L115 | CD2-HD2 | 23.26 | 0.327 | V174 | CG2-HG2 | 21.07 | 0.641 |
| I117 | CD1-HD1 | 16.60 | 0.858 | A177 | CB-HB | 19.53 | 1.416 |
| I117 | CG2-HG2 | 19.61 | 0.760 | V178 | CG1-HG1 | 24.14 | 0.735 |
| M119 | CE-HE | 18.00 | 1.828 | V178 | CG2-HG2 | 21.76 | 0.424 |
| A122 | CB-HB | 20.57 | 1.428 | V180 | CG1-HG1 | 22.09 | 1.007 |
| I123 | CD1-HD1 | 9.14 | 0.750 | V180 | CG2-HG2 | 21.55 | 0.824 |
| I123 | CG2-HG2 | 16.59 | 0.799 | L181 | CD1-HD1 | 23.17 | 0.799 |
| M124 | CE-HE | 17.05 | 2.025 | L181 | CD2-HD2 | 27.17 | 0.630 |
| I128 | CD1-HD1 | 14.36 | 0.546 | I182 | CD1-HD1 | 13.49 | 0.301 |
| I128 | CG2-HG2 | 18.68 | 0.631 | I182 | CG2-HG2 | 16.77 | 0.718 |
| I129 | CD1-HD1 | 14.18 | 0.746 | L185 | CD1-HD1 | 22.10 | 0.724 |
| I129 | CG2-HG2 | 17.55 | 0.904 | L185 | CD2-HD2 | 26.49 | 0.569 |
| L130 | CD1-HD1 | 25.57 | 0.647 | A187 | CB-HB | 17.83 | 1.519 |
| L130 | CD2-HD2 | 24.09 | 0.549 | T191 | CG2-HG2 | 21.11 | 1.152 |
| A132 | CB-HB | 25.38 | 1.374 | V192 | CG1-HG1 | 22.64 | 0.708 |
| V136 | CG1-HG1 | 21.43 | 0.337 | V192 | CG2-HG2 | 22.74 | 0.650 |
| V136 | CG2-HG2 | 20.44 | 0.361 | V194 | CG1-HG1 | 22.07 | 0.830 |
| I137 | CD1-HD1 | 13.45 | 0.691 | V194 | CG2-HG2 | 20.31 | 0.773 |
| I137 | CG2-HG2 | 16.73 | 0.719 | T197 | CG2-HG2 | 23.82 | 1.406 |
| L141 | CD1-HD1 | 25.18 | 0.657 | L198 | CD1-HD1 | 24.55 | 0.575 |
| L141 | CD2-HD2 | 26.62 | 0.333 | L198 | CD2-HD2 | 23.96 | 0.422 |
| T143 | CG2-HG2 | 22.46 | 1.059 | A202 | CB-HB | 20.55 | 1.031 |
| L144 | CD1-HD1 | 23.68 | 0.799 | | | | |

Table S4. Chemical shifts of Ud NS1 methyl atoms.

| Residues | Atom name | $^{13}C$ | $^{1}H$ | Residues | Atom name | $^{13}C$ | $^{1}H$ |
|---|---|---|---|---|---|---|---|
| T84 | CG2-HG2 | 21.59 | 1.258 | I145 | CD1-HD1 | 12.82 | 0.867 |
| A86 | CB-HB | 19.43 | 1.317 | I145 | CG2-HG2 | 17.16 | 0.970 |
| I90 | CD1-HD1 | 9.83 | 0.436 | L146 | CD1-HD1 | 22.89 | 1.195 |
| I90 | CG2-HG2 | 18.18 | 0.592 | L146 | CD2-HD2 | 26.13 | 0.995 |
| T91 | CG2-HG2 | 20.73 | 0.856 | L147 | CD1-HD1 | 28.92 | 0.675 |
| M93 | CE-HE | 18.37 | 1.980 | L147 | CD2-HD2 | 24.14 | 0.536 |
| T94 | CG2-HG2 | 21.94 | 1.273 | A149 | CB-HB | 20.59 | 0.768 |
| I95 | CD1-HD1 | 12.70 | 0.871 | T151 | CG2-HG2 | 23.11 | 1.227 |
| I95 | CG2-HG2 | 17.44 | 0.917 | A155 | CB-HB | 19.34 | 1.360 |
| L98 | CD1-HD1 | 25.55 | 0.832 | I156 | CD1-HD1 | 15.09 | 0.648 |
| L98 | CD2-HD2 | 25.56 | 0.833 | I156 | CG2-HG2 | 16.51 | -0.803 |
| M104 | CE-HE | 17.61 | 1.588 | V157 | CG1-HG1 | 22.97 | 0.814 |
| L105 | CD1-HD1 | 24.53 | 0.954 | V157 | CG2-HG2 | 19.17 | 0.495 |
| L105 | CD2-HD2 | 23.93 | 0.908 | I160 | CD1-HD1 | 14.56 | 0.675 |
| M106 | CE-HE | N.A. | N.A. | I160 | CG2-HG2 | 18.72 | 0.641 |
| V111 | CG1-HG1 | 22.38 | 0.975 | L163 | CD1-HD1 | 26.25 | 0.884 |
| V111 | CG2-HG2 | 22.11 | 0.942 | L163 | CD2-HD2 | 24.12 | 0.970 |
| L115 | CD1-HD1 | 27.23 | 0.487 | T170 | CG2-HG2 | 21.41 | 1.311 |
| L115 | CD2-HD2 | 23.42 | 0.328 | I171 | CD1-HD1 | 14.47 | 1.273 |
| I117 | CD1-HD1 | 16.69 | 0.880 | I171 | CG2-HG2 | 16.74 | 1.073 |
| I117 | CG2-HG2 | 19.40 | 0.787 | V174 | CG1-HG1 | 24.31 | 0.844 |
| I119 | CD1-HD1 | 14.36 | 0.765 | V174 | CG2-HG2 | 20.83 | 0.649 |
| I119 | CG2-HG2 | 17.04 | 0.759 | A177 | CB-HB | 19.55 | 1.385 |
| A122 | CB-HB | 20.74 | 1.418 | I178 | CD1-HD1 | 13.93 | 0.432 |
| I123 | CD1-HD1 | 9.39 | 0.745 | I178 | CG2-HG2 | 17.06 | 0.114 |
| I123 | CG2-HG2 | 16.74 | 0.796 | V180 | CG1-HG1 | 22.39 | 1.008 |
| M124 | CE-HE | 17.20 | 2.047 | V180 | CG2-HG2 | 21.71 | 0.854 |
| I128 | CD1-HD1 | 14.19 | 0.537 | L181 | CD1-HD1 | 24.24 | 0.754 |
| I128 | CG2-HG2 | 19.08 | 0.633 | L181 | CD2-HD2 | 27.91 | 0.617 |
| M129 | CE-HE | 16.49 | 1.933 | I182 | CD1-HD1 | 13.58 | 0.419 |
| L130 | CD1-HD1 | 25.37 | 0.619 | I182 | CG2-HG2 | 16.83 | 0.760 |
| L130 | CD2-HD2 | 24.10 | 0.546 | L185 | CD1-HD1 | 22.49 | 0.741 |
| A132 | CB-HB | 25.45 | 1.422 | L185 | CD2-HD2 | 26.68 | 0.558 |
| V136 | CG1-HG1 | 21.69 | 0.271 | T191 | CG2-HG2 | 21.20 | 1.152 |
| V136 | CG2-HG2 | 20.87 | 0.222 | V192 | CG1-HG1 | 22.72 | 0.726 |
| I137 | CD1-HD1 | 13.53 | 0.693 | V192 | CG2-HG2 | 22.92 | 0.678 |
| I137 | CG2-HG2 | 16.87 | 0.715 | V194 | CG1-HG1 | 22.21 | 0.816 |
| L141 | CD1-HD1 | 25.18 | 0.639 | V194 | CG2-HG2 | 20.35 | 0.791 |
| L141 | CD2-HD2 | 27.15 | 0.003 | T197 | CG2-HG2 | 23.63 | 1.463 |
| T143 | CG2-HG2 | 22.68 | 1.071 | L198 | CD1-HD1 | 25.19 | 0.682 |
| L144 | CD1-HD1 | 23.97 | 0.842 | L198 | CD2-HD2 | 23.06 | 0.293 |
| L144 | CD2-HD2 | 26.19 | 0.559 | A202 | CB-HB | 20.70 | 0.986 |

Table S5. Chemical shifts of Ud NS1 methyl atoms.

| Residues | Atom name | $^{13}C$ | $^{1}H$ | Residues | Atom name | $^{13}C$ | $^{1}H$ |
|---|---|---|---|---|---|---|---|
| M84 | CE-HE | N.A. | N.A. | I145 | CG2-HG2 | 17.77 | 0.939 |
| A86 | CB-HB | 19.47 | 1.317 | L146 | CD1-HD1 | 22.54 | 1.206 |
| L90 | CD1-HD1 | 26.02 | 0.576 | L146 | CD2-HD2 | 26.25 | 1.002 |
| L90 | CD2-HD2 | 24.05 | 0.558 | L147 | CD1-HD1 | 28.43 | 0.710 |
| T91 | CG2-HG2 | 20.93 | 0.802 | L147 | CD2-HD2 | 24.18 | 0.529 |
| M93 | CE-HE | 18.07 | 1.925 | A149 | CB-HB | 20.57 | 0.757 |
| T94 | CG2-HG2 | 22.01 | 1.253 | T151 | CG2-HG2 | 22.99 | 1.247 |
| L95 | CD1-HD1 | 24.27 | 0.913 | A155 | CB-HB | 19.23 | 1.371 |
| L95 | CD2-HD2 | 24.16 | 0.886 | I156 | CD1-HD1 | 15.46 | 0.673 |
| M98 | CE-HE | 17.00 | 1.996 | I156 | CG2-HG2 | 16.74 | -0.820 |
| M104 | CE-HE | 17.66 | 1.609 | V157 | CG1-HG1 | 22.68 | 0.823 |
| L105 | CD1-HD1 | 24.46 | 0.964 | V157 | CG2-HG2 | 18.82 | 0.468 |
| L105 | CD2-HD2 | 24.31 | 0.939 | I160 | CD1-HD1 | 14.36 | 0.673 |
| M106 | CE-HE | N.A. | N.A. | I160 | CG2-HG2 | 18.74 | 0.667 |
| V111 | CG1-HG1 | 21.52 | 0.925 | L163 | CD1-HD1 | 25.85 | 0.912 |
| V111 | CG2-HG2 | 21.52 | 0.822 | L163 | CD2-HD2 | 23.31 | 0.870 |
| A112 | CB-HB | 20.18 | 1.202 | L166 | CD1-HD1 | N.A. | N.A. |
| L115 | CD1-HD1 | 27.05 | 0.544 | L166 | CD2-HD2 | N.A. | N.A. |
| L115 | CD2-HD2 | 23.15 | 0.247 | T170 | CG2-HG2 | 21.50 | 1.311 |
| I117 | CD1-HD1 | 15.95 | 0.861 | V174 | CG1-HG1 | 26.53 | 1.013 |
| I117 | CG2-HG2 | 19.57 | 0.769 | V174 | CG2-HG2 | 21.73 | 0.774 |
| M119 | CE-HE | 18.03 | 1.833 | A177 | CB-HB | 19.36 | 1.435 |
| A122 | CB-HB | 20.50 | 1.419 | I178 | CD1-HD1 | 14.56 | 0.526 |
| I123 | CD1-HD1 | 9.32 | 0.746 | I178 | CG2-HG2 | 17.17 | 0.141 |
| I123 | CG2-HG2 | 16.76 | 0.806 | V180 | CG1-HG1 | 22.63 | 1.012 |
| M124 | CE-HE | 17.12 | 2.050 | V180 | CG2-HG2 | 21.52 | 0.822 |
| T127 | CG2-HG2 | 21.52 | 1.065 | L181 | CD1-HD1 | 23.42 | 0.803 |
| I128 | CD1-HD1 | 14.38 | 0.545 | L181 | CD2-HD2 | 27.21 | 0.604 |
| I128 | CG2-HG2 | 18.91 | 0.620 | I182 | CD1-HD1 | 13.22 | 0.079 |
| I129 | CD1-HD1 | 14.51 | 0.738 | I182 | CG2-HG2 | 16.70 | 0.715 |
| I129 | CG2-HG2 | 17.83 | 0.858 | L185 | CD1-HD1 | 22.28 | 0.732 |
| L130 | CD1-HD1 | 25.06 | 0.624 | L185 | CD2-HD2 | 26.59 | 0.546 |
| L130 | CD2-HD2 | 24.43 | 0.550 | A187 | CB-HB | 17.89 | 1.532 |
| A132 | CB-HB | 24.12 | 1.174 | T191 | CG2-HG2 | 21.12 | 1.172 |
| V136 | CG1-HG1 | 21.31 | 0.322 | V192 | CG1-HG1 | 22.82 | 0.714 |
| V136 | CG2-HG2 | 20.57 | 0.348 | V192 | CG2-HG2 | 22.89 | 0.720 |
| I137 | CD1-HD1 | 13.71 | 0.689 | V194 | CG1-HG1 | 21.99 | 0.803 |
| I137 | CG2-HG2 | 16.91 | 0.702 | V194 | CG2-HG2 | 19.02 | 0.905 |
| L141 | CD1-HD1 | 25.45 | 0.674 | T195 | CG2-HG2 | 22.56 | 1.570 |
| L141 | CD2-HD2 | 26.47 | 0.218 | T197 | CG2-HG2 | 24.82 | 1.316 |
| T143 | CG2-HG2 | 22.61 | 1.094 | I198 | CD1-HD1 | 14.32 | 0.859 |
| L144 | CD1-HD1 | 24.15 | 0.863 | I198 | CG2-HG2 | 17.77 | 0.811 |
| L144 | CD2-HD2 | 26.39 | 0.585 | A202 | CB-HB | 21.73 | 1.013 |
| I145 | CD1-HD1 | 12.68 | 0.826 | | | | |

Supplementary Figure 1

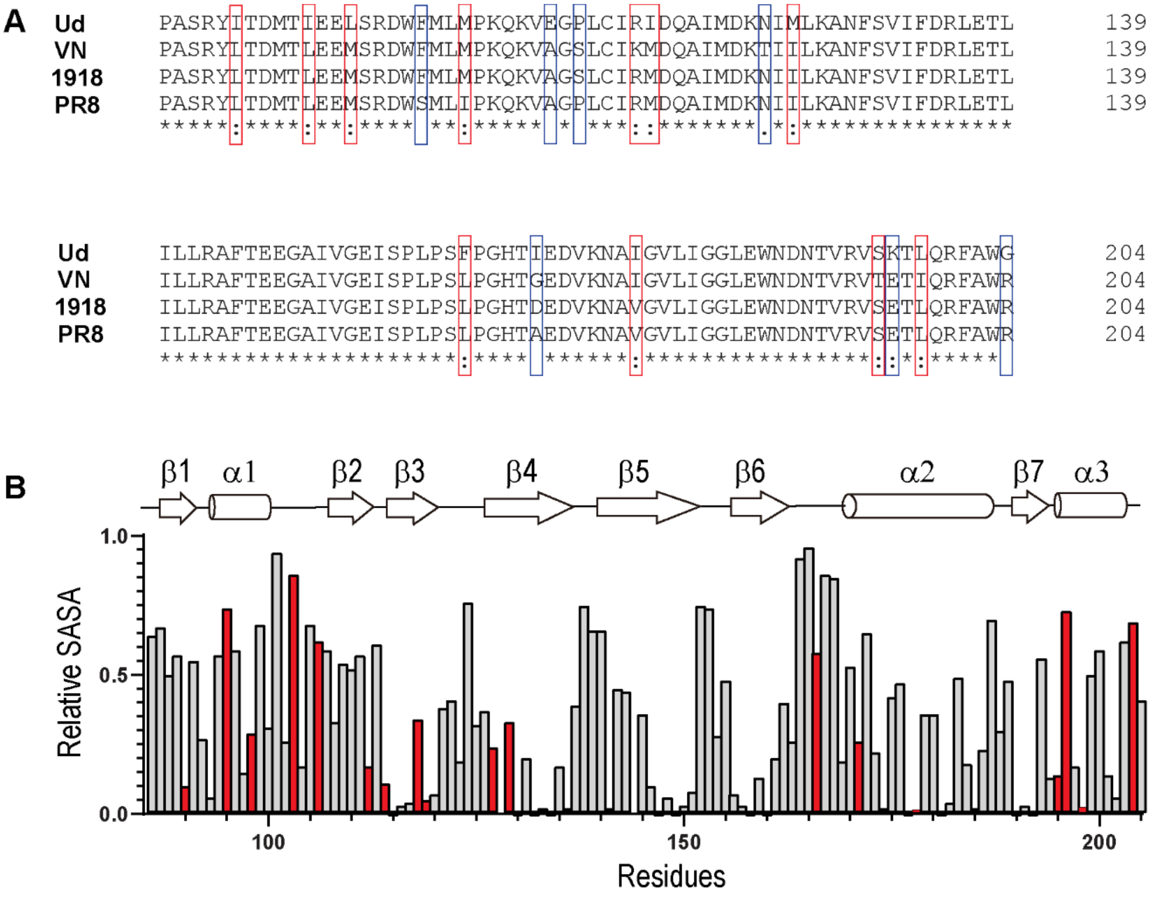

**Supplementary Figure 1**. (A) Amino acids sequence alignment of NS1 ED proteins (residues 85 – 205). Sequence alignment was performed by the Clustal Omega webserver (https://www.ebi.ac.uk/Tools/msa/clustalo/). Positions for conservative and non-conservative mutations are highlighted by red and blue boxes, respectively. (B) Relative solvent accessible surface area (SASA) for individual residues of NS1. Positions for strain-specific mutations are highlighted in red. The relative SASA was calculated for 1918 NS1 (PDB ID: 6U28, molecule A) using Pymol.

Supplementary Figure 2

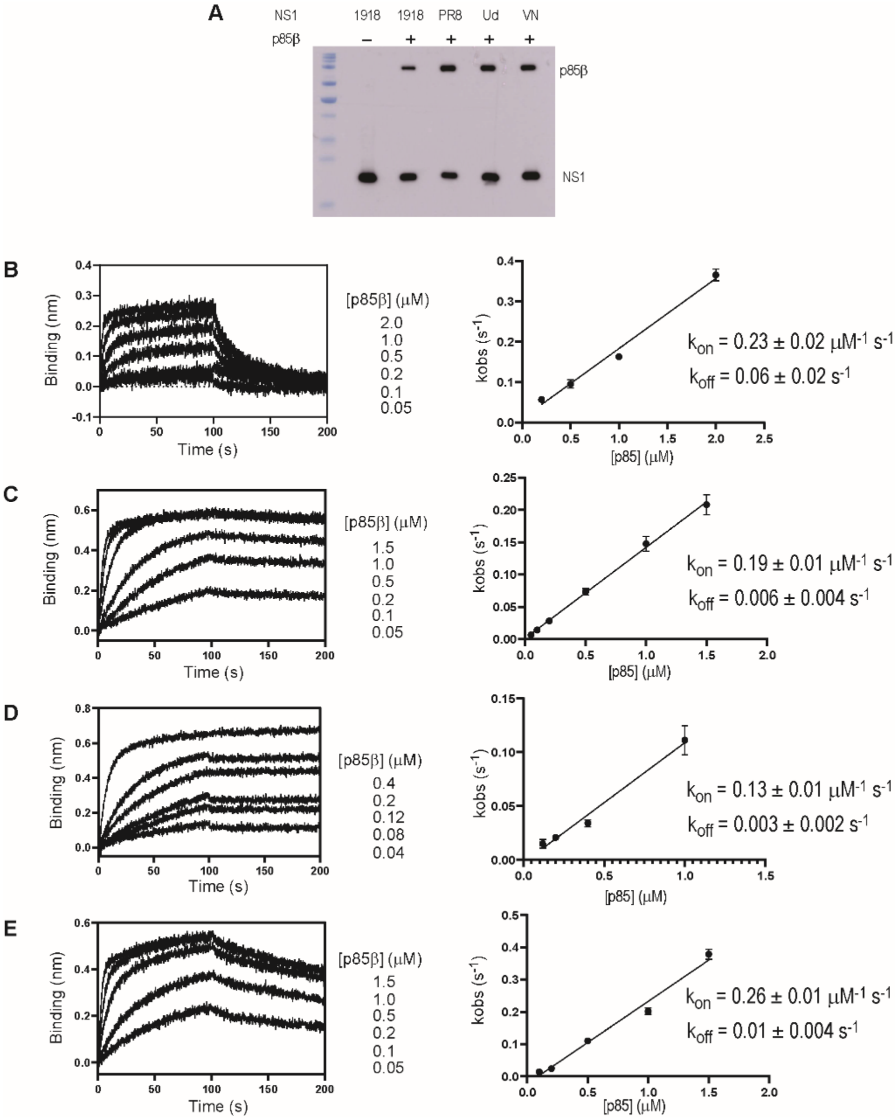

**Supplementary Figure 2.** Interactions between NS1s and p85β. (A) In vitro co-immunoprecipitation of $His_6$-tagged PI3K by biotin-$His_6$-tagged NS1s. 3 μg of full-length PI3K (p85β and p110 subunits) was mixed with 3 μg of individual NS1s. The full-length PI3K was purchased from ThermoFisher Scientific (cat. no. A31084). Only p85β has the N-terminal $His_6$-tag for visualization using the anti-His antibody. Representative BLI sensorgrams and plots of $k_{obs}$ vs [p85] for (B) 1918, (B) Ud, (C) PR8, and (D) VN NS1s. Fit parameters are shown in the plot. Fit parameters and uncertainties are averages and standard deviations of three repeated measurements.

Supplementary Figure 3

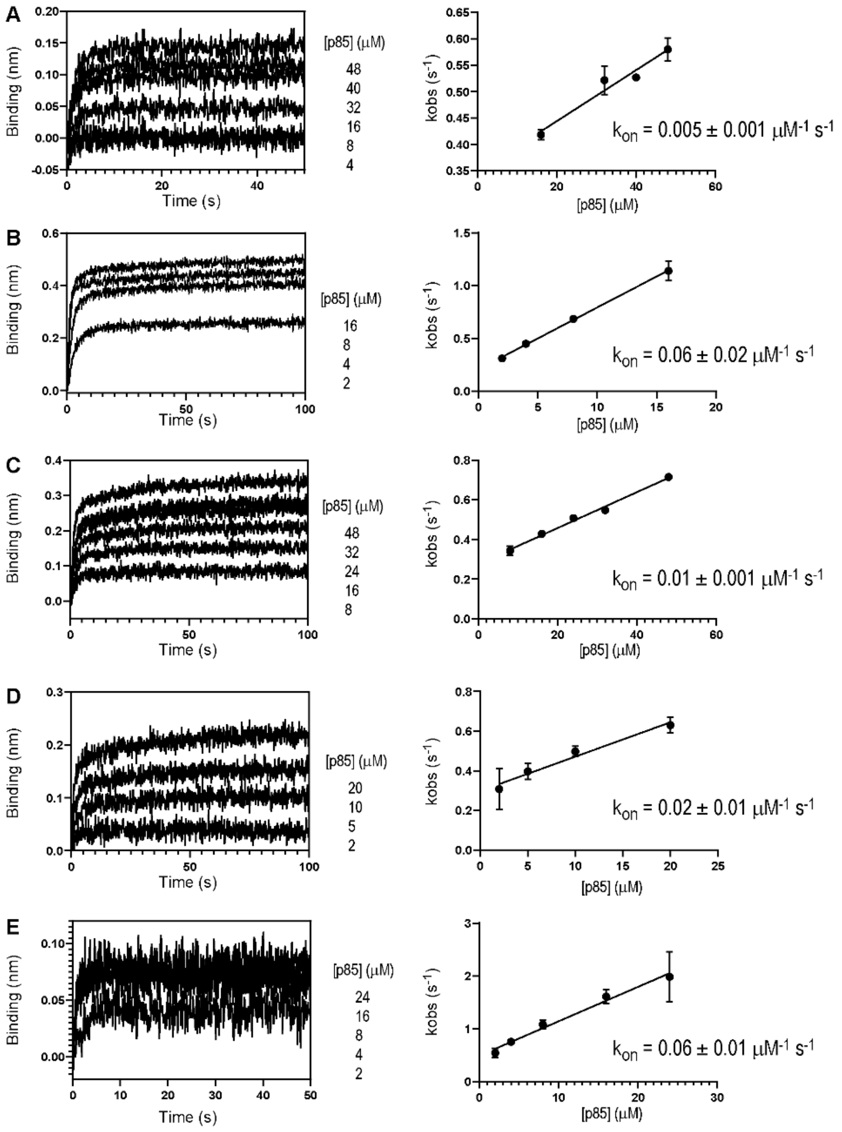

**Supplementary Figure 3.** Representative BLI sensorgrams (association phases) and plots of $k_{obs}$ vs. [p85] for 1918 NS1 (A) Y89F, (B) L95A, (C) M98A, (D) I145A, and (E) L146A. Fit parameters and uncertainties are averages and standard deviations of three repeated measurements.

Supplementary Figure 3 (continued)

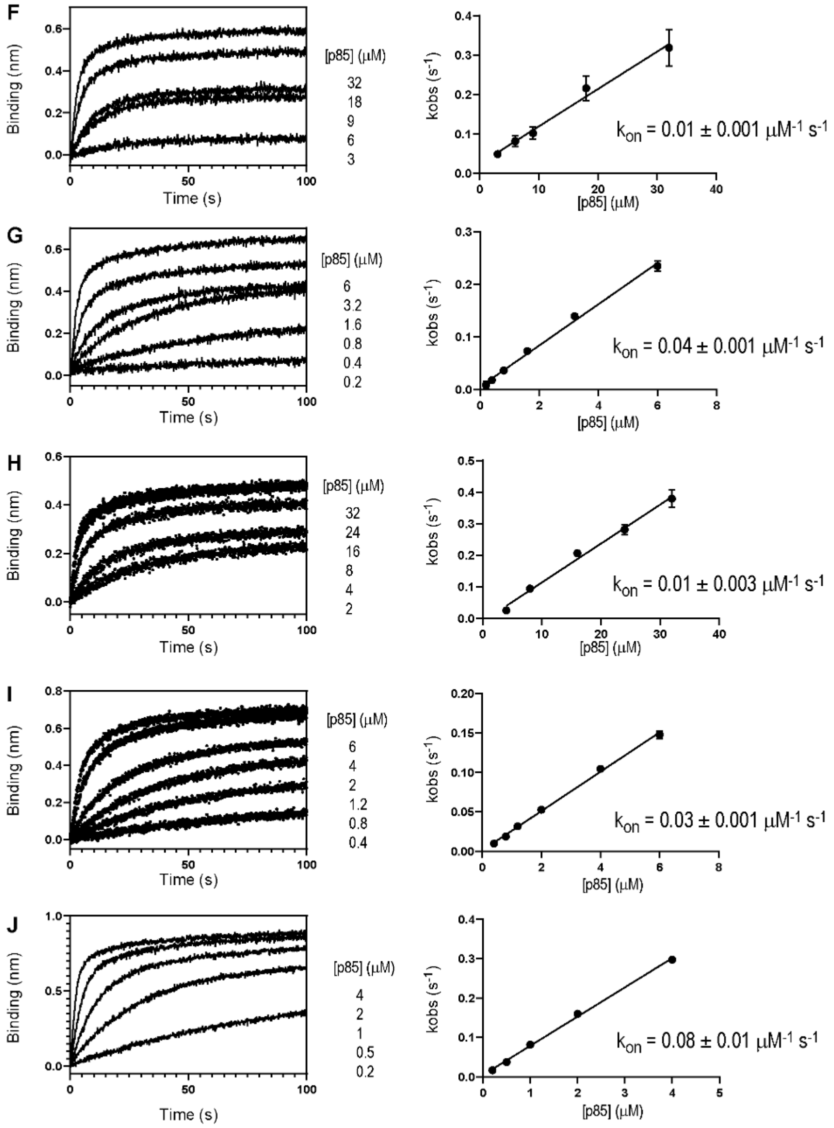

**Supplementary Figure 3 (continued).** Representative BLI sensorgrams (association phases) and plots of kobs vs. [p85] for PR8 NS1 (F) Y89F, (G) L95A, (H) M98A, (I) I145A, and (J) L146A. Fit parameters and uncertainties are averages and standard deviations of three repeated measurements.

Supplementary Figure 3 (continued)

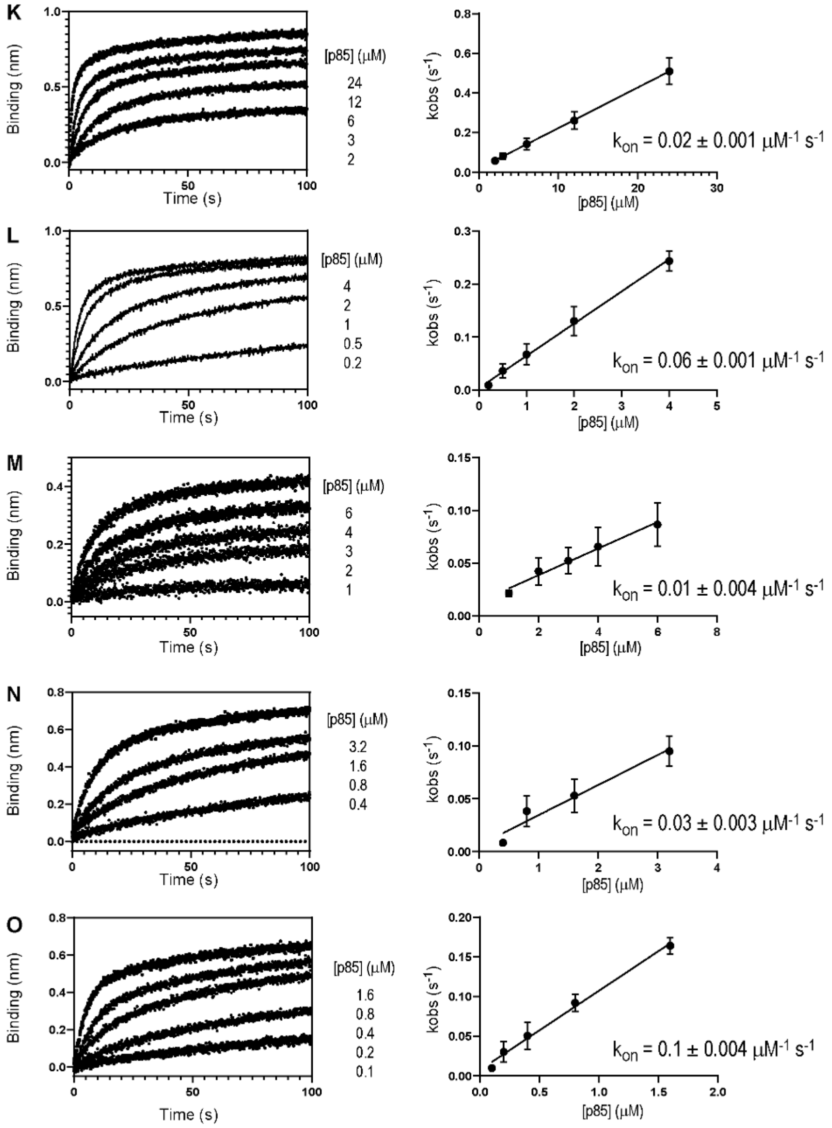

**Supplementary Figure 3 (continued).** Representative BLI sensorgrams (association phases) and plots of kobs vs. [p85] for and Ud NS1 (K) Y89F, (L) I95A, (M) L98A, (N) I145A, and (O) L146A. Fit parameters and uncertainties are averages and standard deviations of three repeated measurements.

Supplementary Figure 3 (continued)

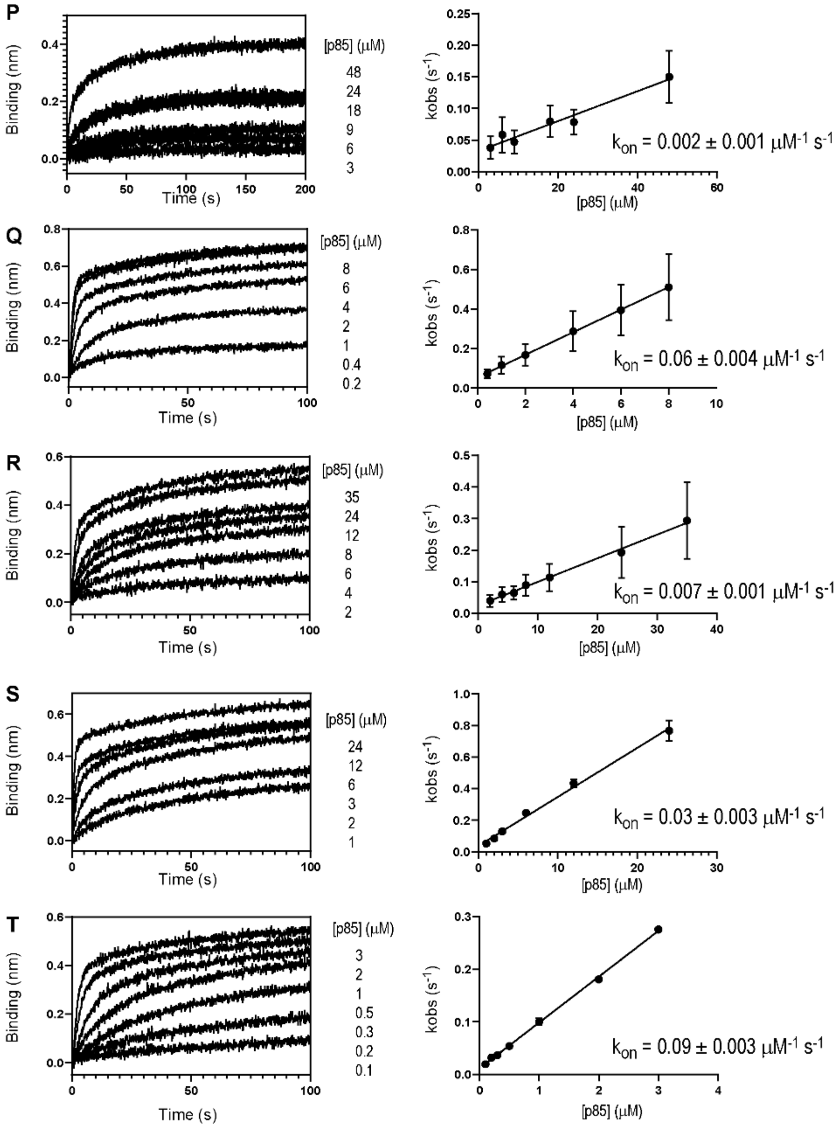

**Supplementary Figure 3 (continued).** Representative BLI sensorgrams (association phases) and plots of kobs vs. [p85] for VN NS1 (P) Y89F, (Q) L95A, (R) M98A, (S) I145A, and (T) L146A. Fit parameters and uncertainties are averages and standard deviations of three repeated measurements.

Supplementary Figure 4

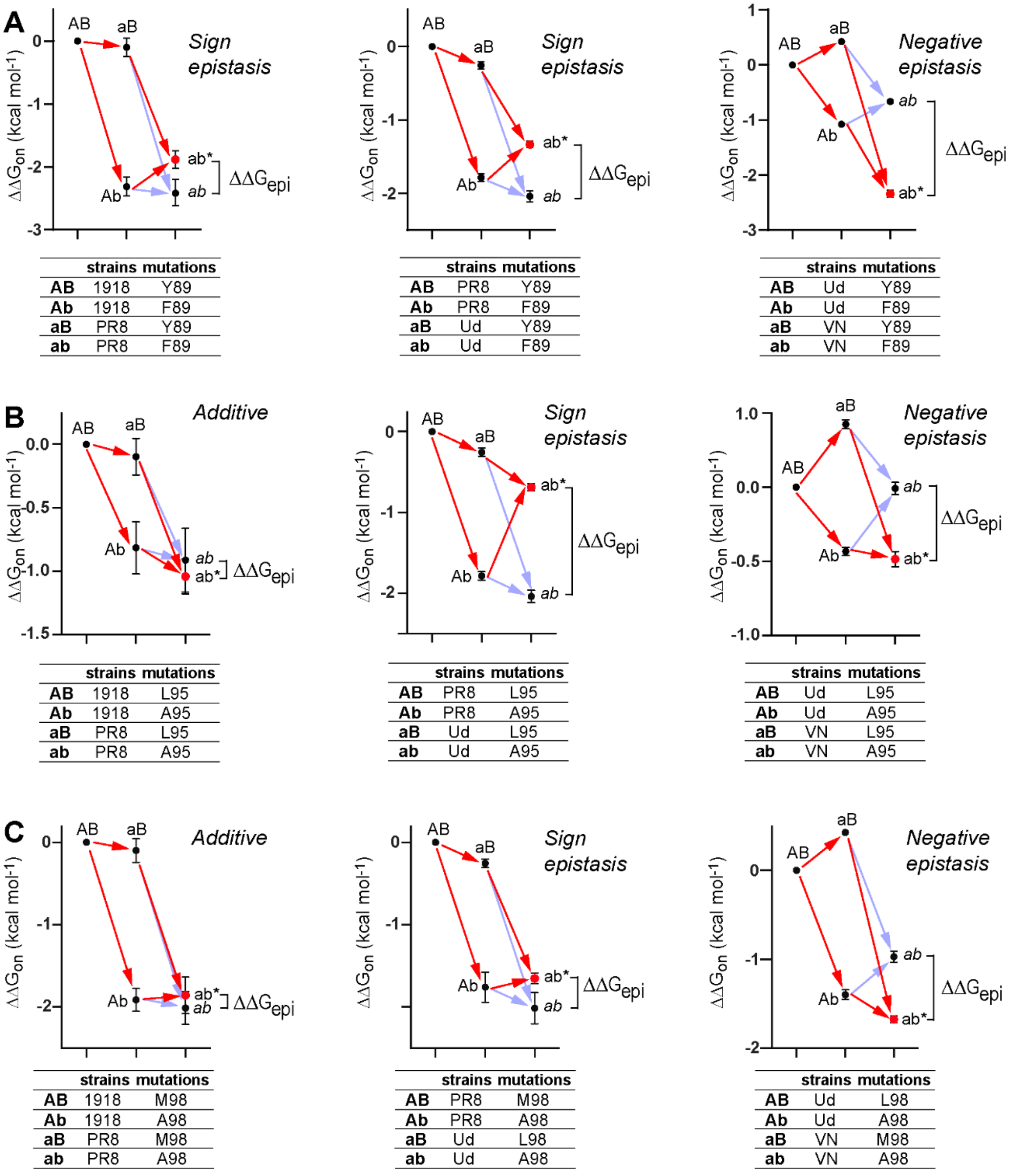

| strains | | mutations |
|---|---|---|
| **AB** | 1918 | Y89 |
| **Ab** | 1918 | F89 |
| **aB** | PR8 | Y89 |
| **ab** | PR8 | F89 |

| strains | | mutations |
|---|---|---|
| **AB** | PR8 | Y89 |
| **Ab** | PR8 | F89 |
| **aB** | Ud | Y89 |
| **ab** | Ud | F89 |

| strains | | mutations |
|---|---|---|
| **AB** | Ud | Y89 |
| **Ab** | Ud | F89 |
| **aB** | VN | Y89 |
| **ab** | VN | F89 |

| strains | | mutations |
|---|---|---|
| **AB** | 1918 | L95 |
| **Ab** | 1918 | A95 |
| **aB** | PR8 | L95 |
| **ab** | PR8 | A95 |

| strains | | mutations |
|---|---|---|
| **AB** | PR8 | L95 |
| **Ab** | PR8 | A95 |
| **aB** | Ud | L95 |
| **ab** | Ud | A95 |

| strains | | mutations |
|---|---|---|
| **AB** | Ud | L95 |
| **Ab** | Ud | A95 |
| **aB** | VN | L95 |
| **ab** | VN | A95 |

| strains | | mutations |
|---|---|---|
| **AB** | 1918 | M98 |
| **Ab** | 1918 | A98 |
| **aB** | PR8 | M98 |
| **ab** | PR8 | A98 |

| strains | | mutations |
|---|---|---|
| **AB** | PR8 | M98 |
| **Ab** | PR8 | A98 |
| **aB** | Ud | L98 |
| **ab** | Ud | A98 |

| strains | | mutations |
|---|---|---|
| **AB** | Ud | L98 |
| **Ab** | Ud | A98 |
| **aB** | VN | M98 |
| **ab** | VN | A98 |

**Supplementary Figure 4.** Thermodynamic cycle analyses between strain-specific NS1 mutations and core interface mutations on residues (A) 89, (B) 95, (C) 98, (D) 145, and (E) 146. Faint blue arrows represent the expected additive effects of double mutations. Red arrows represent the experimentally measured mutational effects. The difference between ab* and *ab* defines the pattern of epistasis. $\Delta\Delta G_{epi}$ (ab* – *ab*) corresponds to the strength of epistatic interactions.

Supplementary Figure 4 (continued)

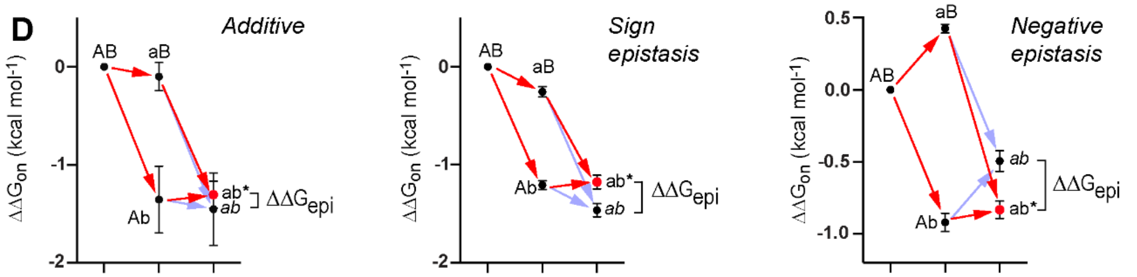

| | strains | mutations |
|---|---|---|
| **AB** | 1918 | I145 |
| **Ab** | 1918 | A145 |
| **aB** | PR8 | I145 |
| **ab** | PR8 | A145 |

| | strains | mutations |
|---|---|---|
| **AB** | PR8 | I145 |
| **Ab** | PR8 | A145 |
| **aB** | Ud | I145 |
| **ab** | Ud | A145 |

| | strains | mutations |
|---|---|---|
| **AB** | Ud | I145 |
| **Ab** | Ud | A145 |
| **aB** | VN | I145 |
| **ab** | VN | A145 |

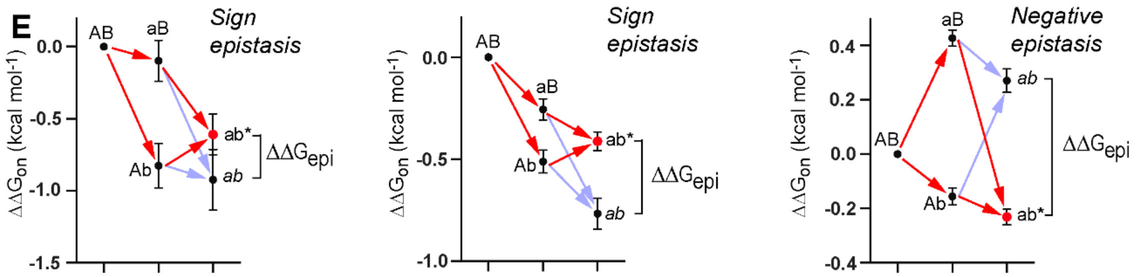

| | strains | mutations |
|---|---|---|
| **AB** | 1918 | L146 |
| **Ab** | 1918 | A146 |
| **aB** | PR8 | L146 |
| **ab** | PR8 | A146 |

| | strains | mutations |
|---|---|---|
| **AB** | PR8 | L146 |
| **Ab** | PR8 | A146 |
| **aB** | Ud | L146 |
| **ab** | Ud | A146 |

| | strains | mutations |
|---|---|---|
| **AB** | Ud | L146 |
| **Ab** | Ud | A146 |
| **aB** | VN | L146 |
| **ab** | VN | A146 |

Supplementary Figure 5

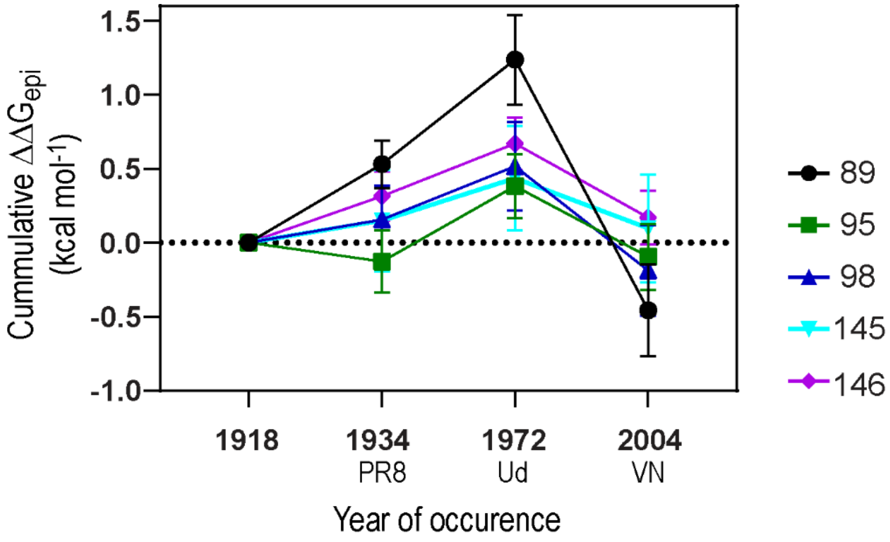

**Supplementary Figure 5.** Cumulative epistatic interactions with respect to 1918 NS1. Cumulative values were calculated by adding $\Delta\Delta G_{epi}$ values from 1918. For example, cumulative $\Delta\Delta G_{epi}$ of Y89F ($\Delta\Delta G_{epi}^{Y89F}$) from 1918 to 2004 (VN) was calculated as follows; $\Delta\Delta G_{epi}^{Y89F}$(2004 – 1918) = $\Delta\Delta G_{epi}^{Y89F}$(1934 – 1918) + $\Delta\Delta G_{epi}^{Y89F}$(1972 – 1934) + $\Delta\Delta G_{epi}^{Y89F}$(2004 – 1972). Individual $\Delta\Delta G_{epi}$(A – B) values are defined in Supplementary Figure 4, where A and B correspond to two different years. For example, $\Delta\Delta G_{epi}^{Y89F}$(1934 – 1918), $\Delta\Delta G_{epi}^{Y89F}$(1972 – 1934), and $\Delta\Delta G_{epi}^{Y89F}$(2004 – 1972) correspond to $\Delta\Delta G_{epi}$ defined in the left, middle, and right panels of Fig S4A.

Supplementary Figure 6A

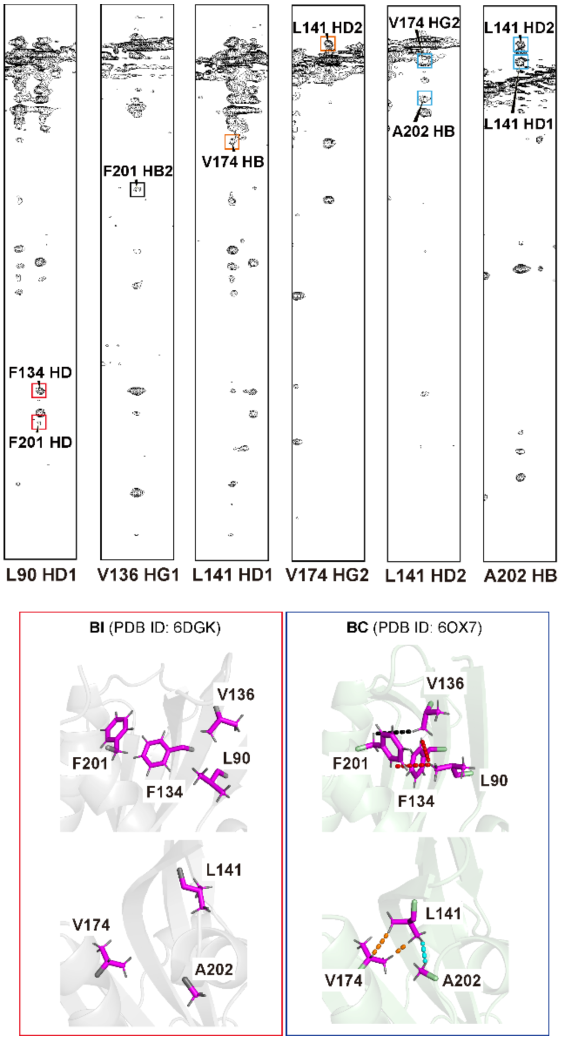

**Supplementary Figure 6A.** NMR NOESY spectra of free 1918 NS1 (upper panels) and crystal structures corresponding to BI and BC conformers of free 1918 NS1. The identified $^{1}$H atom pairs are marked by dotted lines in the BC structures (lower right panel) in the same color as the corresponding NOESY cross-peaks (upper panels). The same $^{1}$H atom pairs are > 5 Å in the BI conformer (gray).

Supplementary Figure 6B

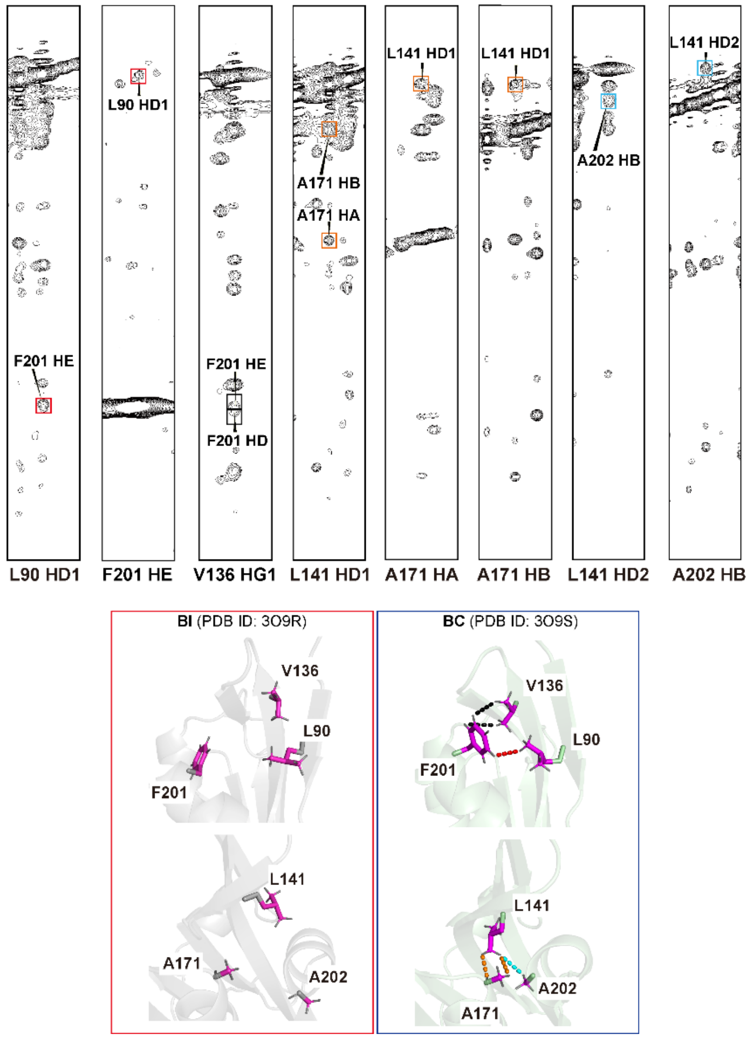

**Supplementary Figure 6B.** NMR NOESY spectra of free PR8 NS1 (upper panels) and crystal structures corresponding to BI and BC conformers of free PR8 NS1. The identified $^{1}$H atom pairs are marked by dotted lines in the BC structures (lower right panel) in the same color as the corresponding NOESY cross-peaks (upper panels). The same $^{1}$H atom pairs are > 5 Å in the BI conformer (gray).

Supplementary Figure 6C

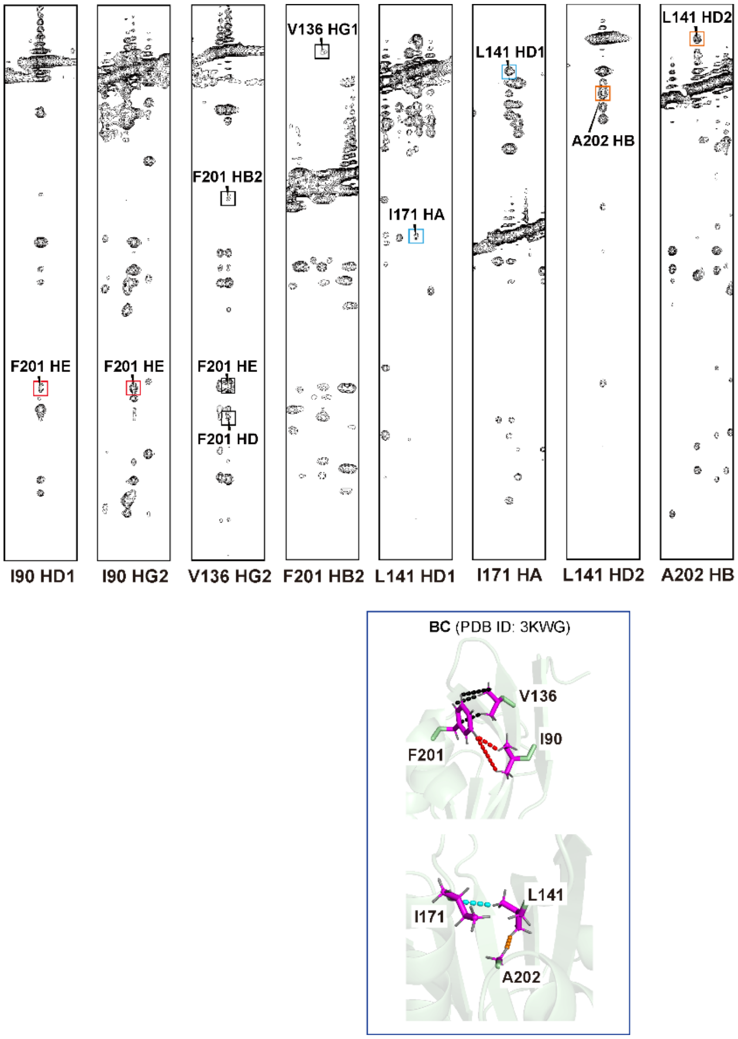

**Supplementary Figure 6C.** NMR NOESY spectra of free Ud NS1 (upper panels) and the crystal structure corresponding to the BC conformer. The identified $^{1}$H atom pairs are marked by dotted lines in the BC structures (lower right panel) in the same color as the corresponding NOESY cross-peaks (upper panels). There is no structure corresponding to the BI conformer of Ud NS1 in PDB.

Supplementary Figure 6D

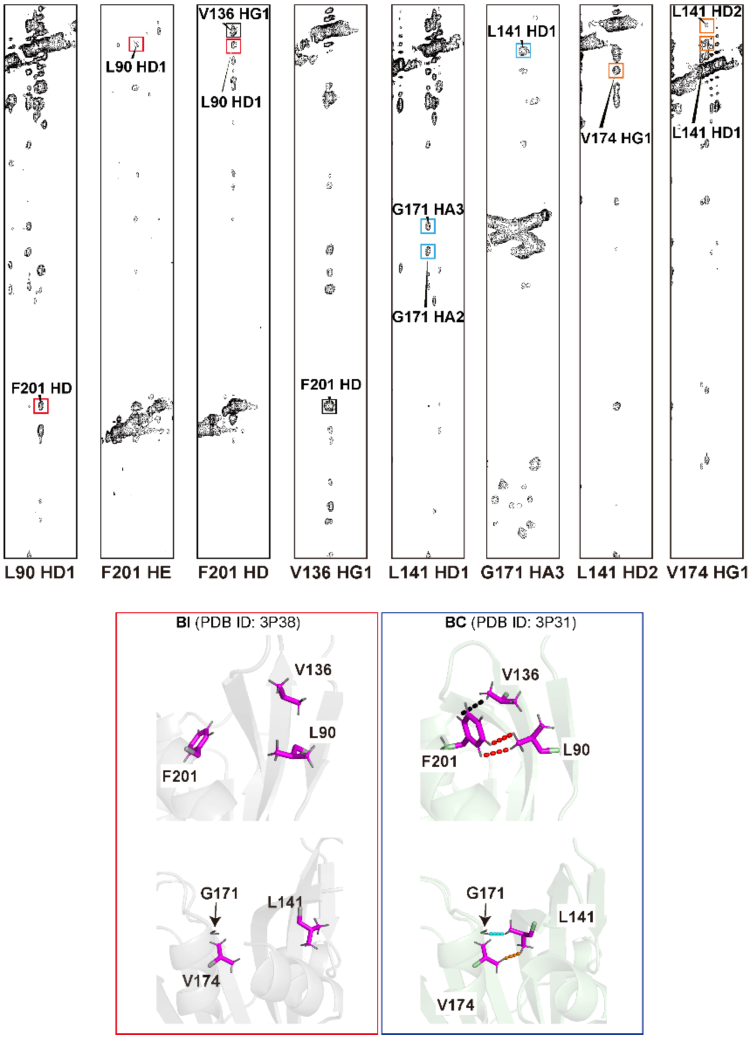

**Supplementary Figure 6D.** NMR NOESY spectra of free VN NS1 (upper panels) and crystal structures corresponding to BI and BC conformers of free VN NS1. The identified $^{1}H$ atom pairs are marked by dotted lines in the BC structures (lower right panel) in the same color as the corresponding NOESY cross-peaks (upper panels). The same $^{1}H$ atom pairs are > 5 Å in the BI conformer (gray).

Supplementary Figure 7A

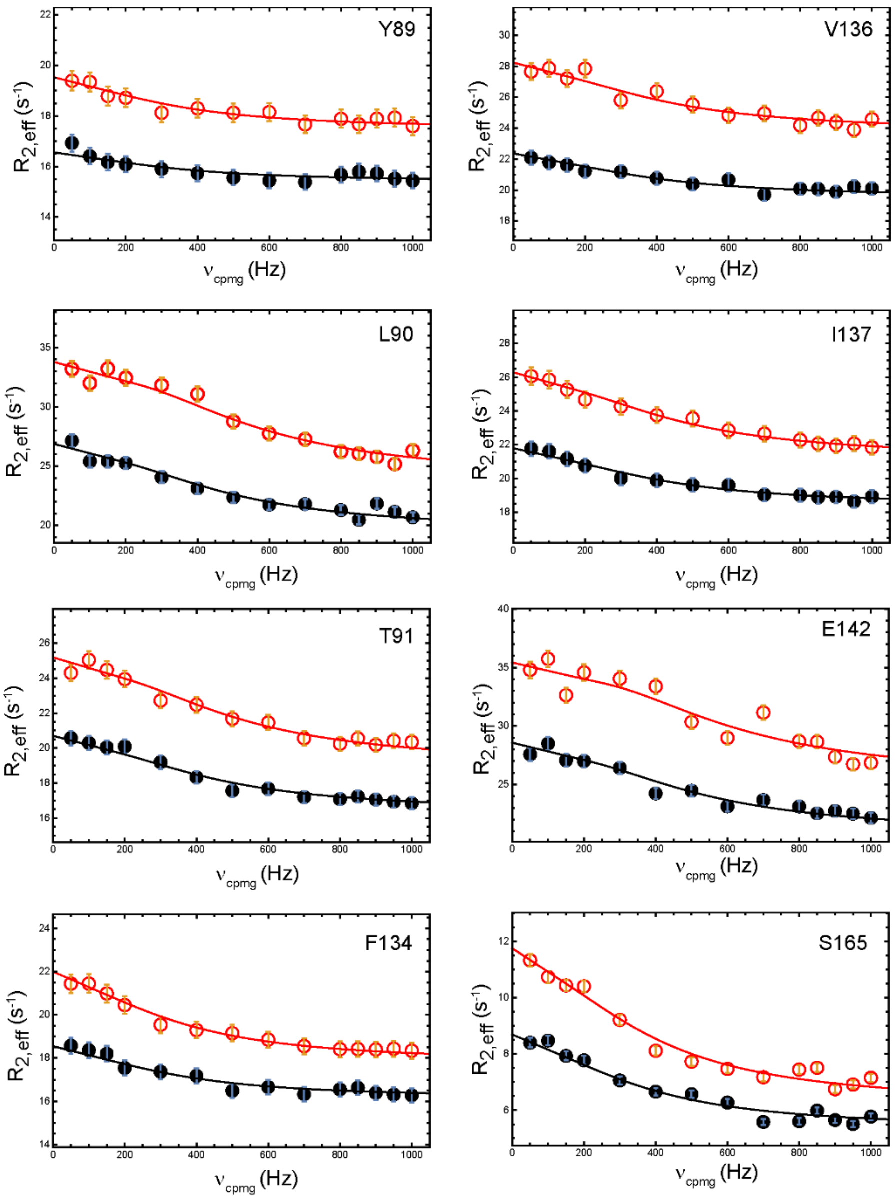

**Supplementary Figure 7A.** NMR $^{15}N$ CPMG-RD data of free PR8 NS1. Data at both 600- (black circles) and 800-MHz (red circles) were globally fit (solid lines) using the Carver-Richards equation with an assumption of a simple two-state exchange model. All residues were fit with a globally constrained $k_{ex}$ value (2200 ± 930 $s^{-1}$). The population of the minor species was derived from slow-exchanging residues (residues 90, 134, 137, 197, and 201).

Supplementary Figure 7A (continued)

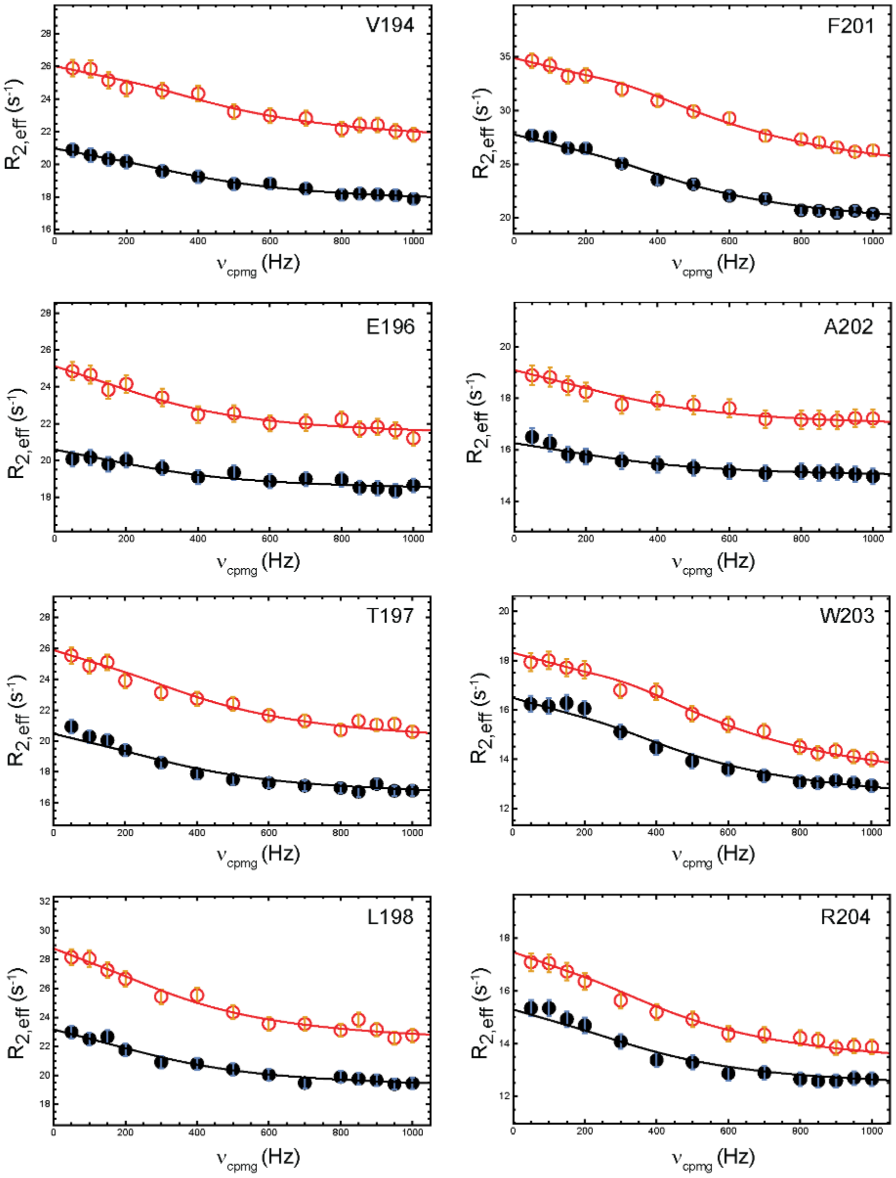

Supplementary Figure 7B

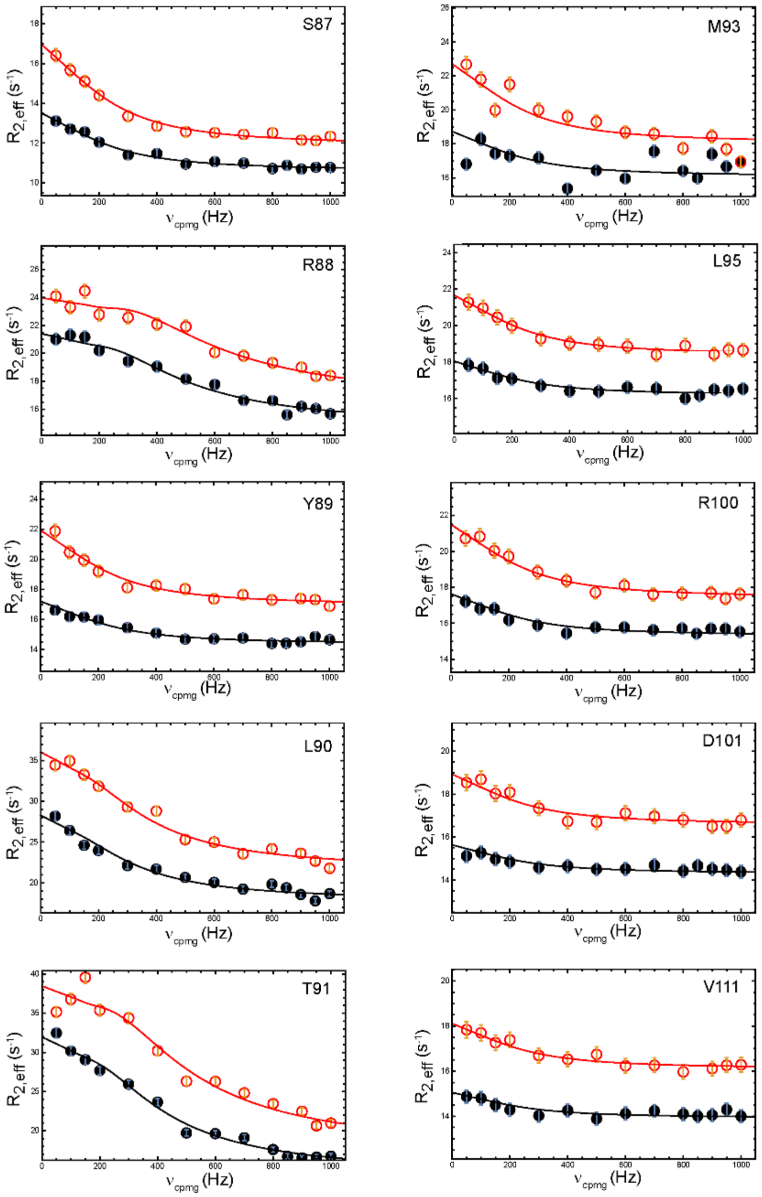

**Supplementary Figure 7B.** NMR $^{15}$N CPMG-RD data of free VN NS1. Data at both 600- (black circles) and 800-MHz (red circles) were globally fit (solid lines) using the Carver-Richards equation with an assumption of a simple two-state exchange model. All residues were fit with a globally constrained $k_{ex}$ value (1500 ± 400 $s^{-1}$). The population of the minor species was derived from slow-exchanging residues (residues 88, 90, 91,127, and 153).

Supplementary Figure 7B (continued)

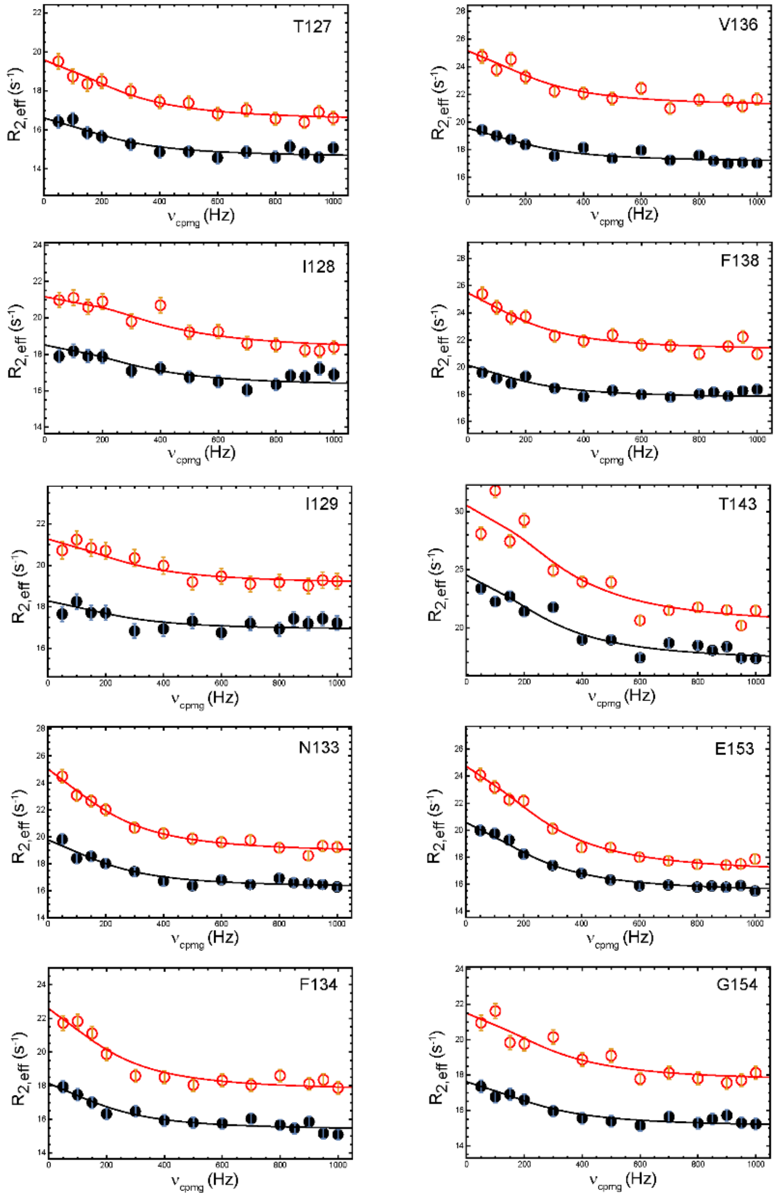

Supplementary Figure 7B (continued)

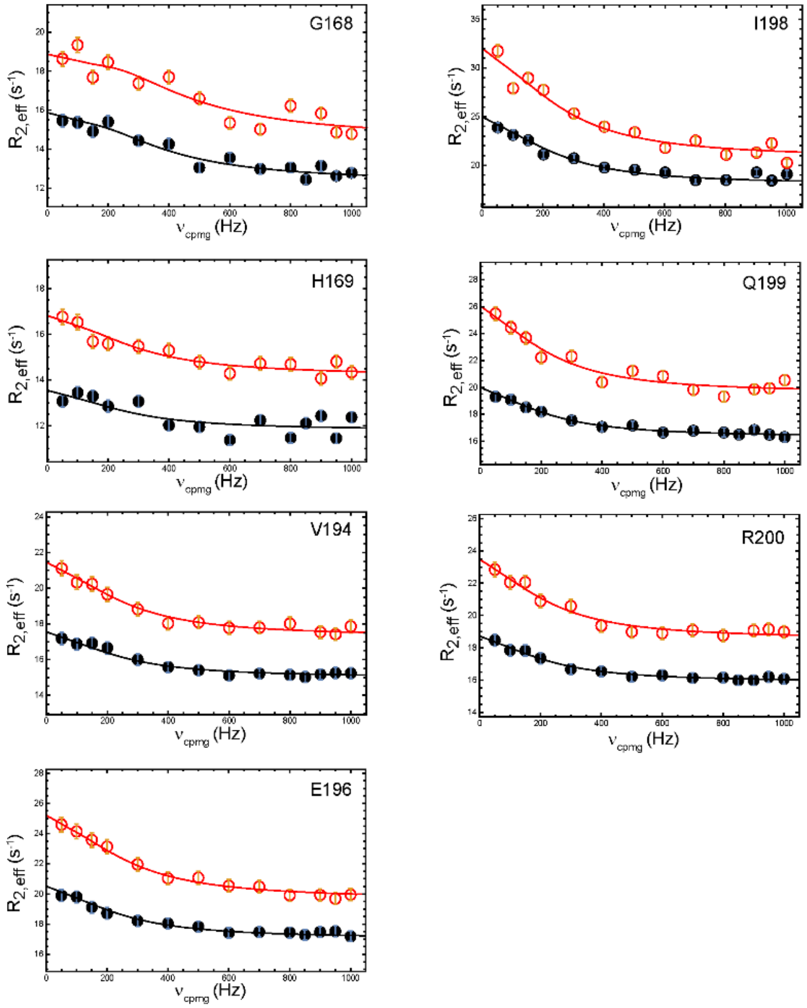

Supplementary Figure 8

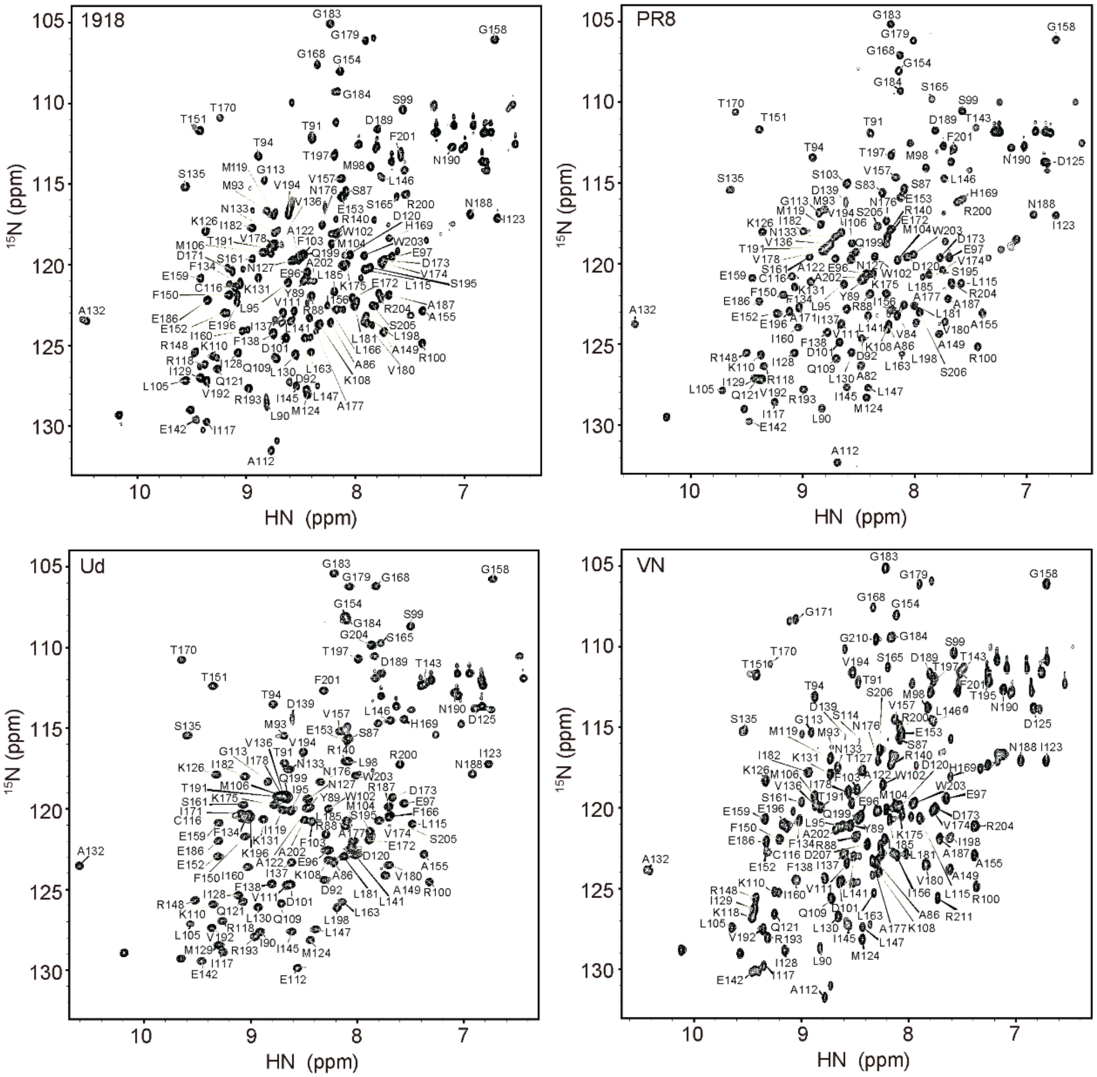

**Supplementary Figure 8.** $^{1}$H-$^{15}$N HSQC spectra of free NS1 proteins. The assignments of backbone resonances are available from the BMRB: 1918 (BMRB accession number: 12032), PR8 (BMRB accession number: 51404), Ud (BMRB accession number: 16376), and VN (BMRB accession number: 51403).

Supplementary Figure 9

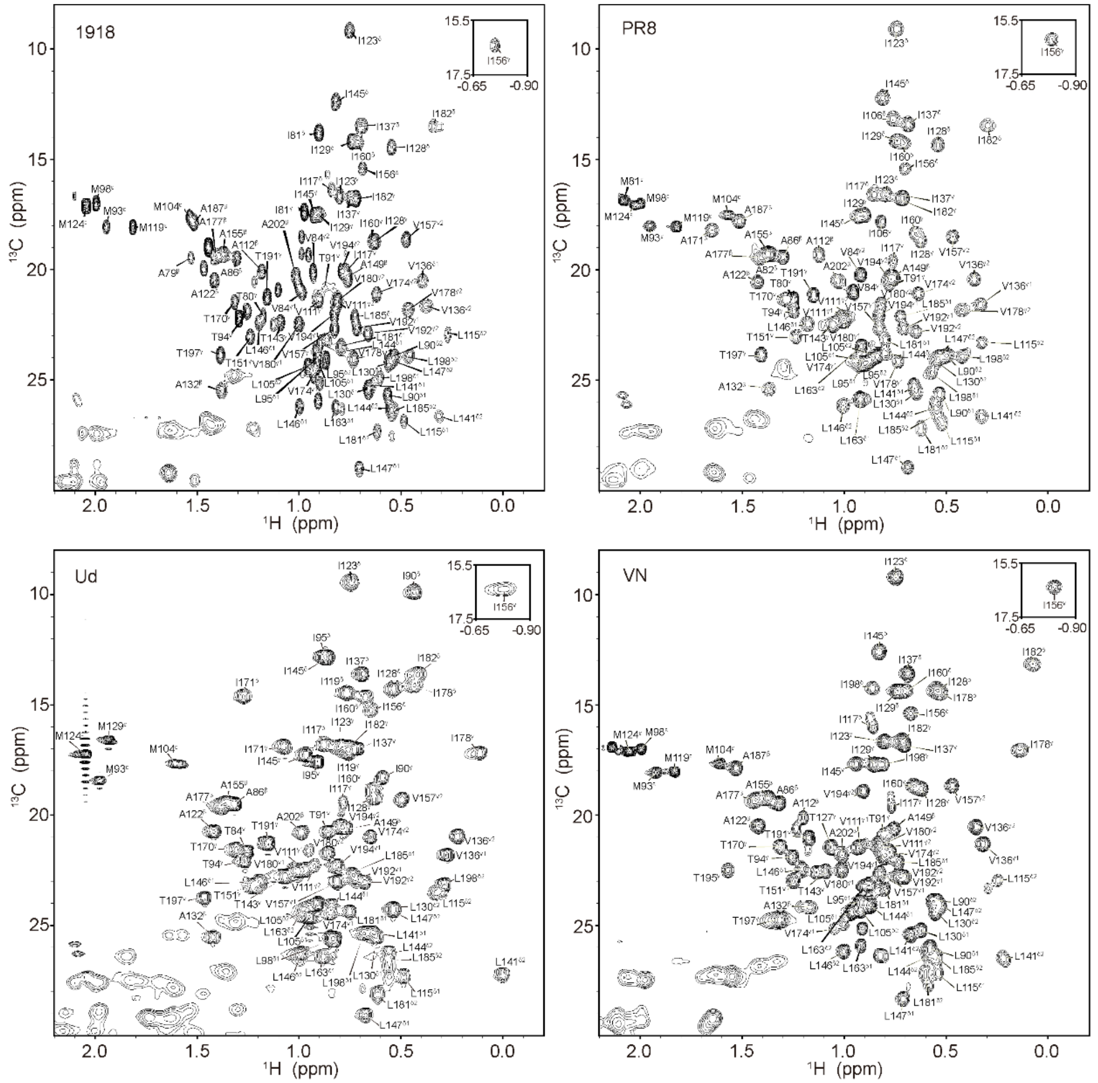

**Supplementary Figure 9.** $^{1}$H-$^{13}$C HSQC spectra for methyl resonances of free NS1 proteins. The assignments of methyl resonances are shown in Supplementary Table S2-S5.

Supplementary Figure 10

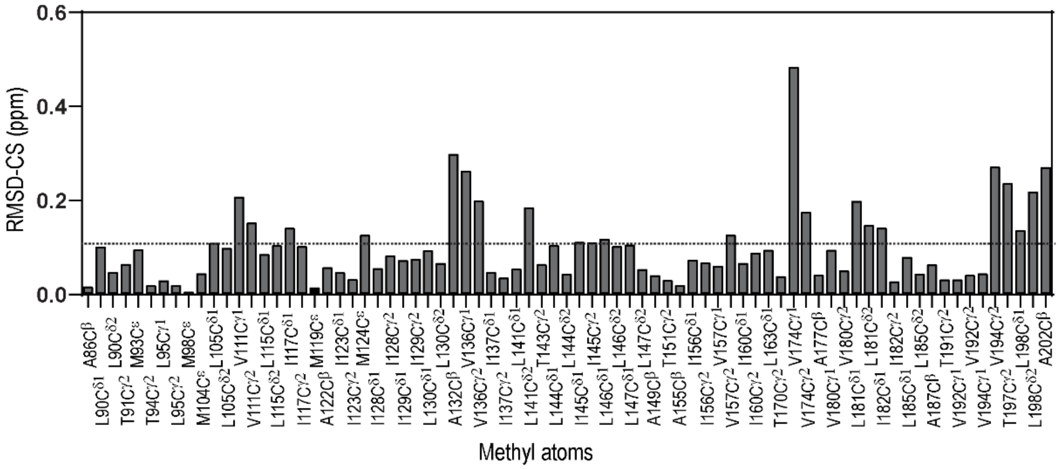

**Supplementary Figure 10.** RMSD of methyl resonances ($^1$H and $^{13}$C) of NS1s. The dotted line represents Q3 of RMSD values.

Supplementary Figure 11

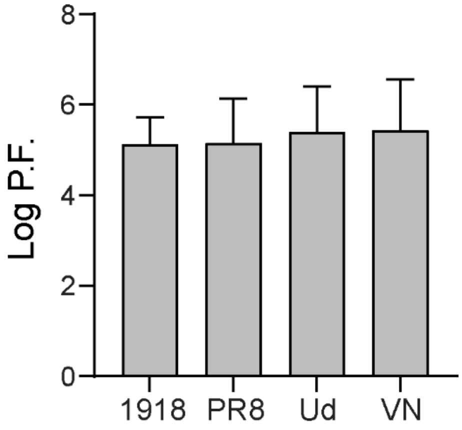

**Supplementary Figure 11.** Average protection factors (P.F.) of backbone amide proton exchange of NS1s in $D_2O$.

Supplementary Figure 12

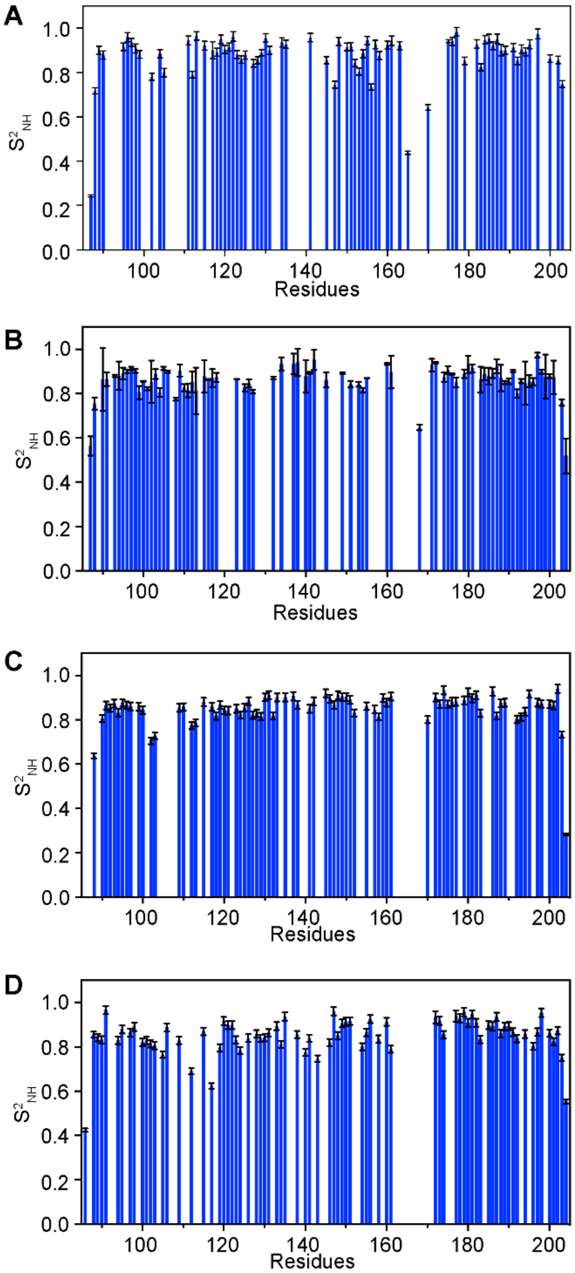

**Supplementary Figure 12.** Order parameters for backbone $^{15}$N-H vectors ($S^2_{NH}$) of (A) 1918, (B) PR8, (C) Ud, and (D) VN NS1s in the free state.

Supplementary Figure 13

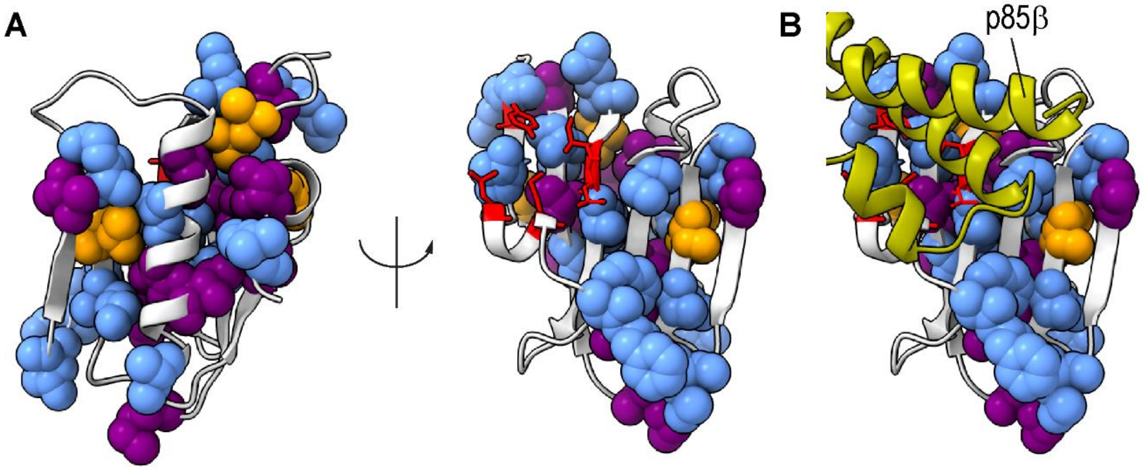

**Supplementary Figure 13.** Structure of NS1 (1918 NS1, PDB ID: 6U28). (A) Residues whose backbone is dynamically variable across NS1s (i.e., $S^2_{NH}$ > Q3) are shown in blue spheres. Residues whose side-chain is conformationally variable across NS1s (i.e., RMSD-CS,$^{13}CH_3$ > Q3) are shown in purple spheres. Residues with both dynamically variable backbone and conformationally variable side-chain are shown in orange. Core interface residues are shown in red sticks. (B) The bound p85β is shown to display the binding interface on NS1.

Supplementary Figure 14

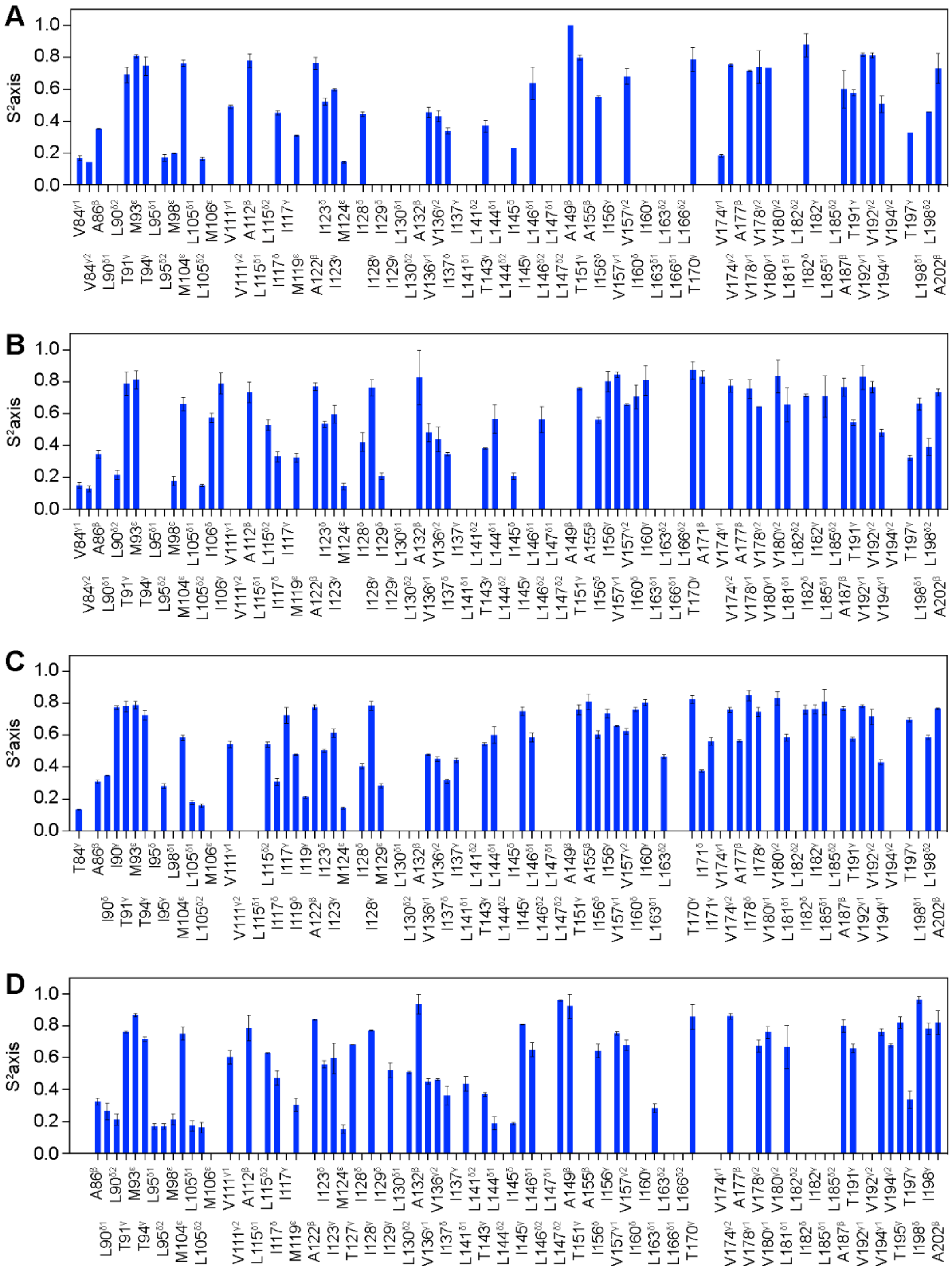

**Supplementary Figure 14.** NMR order parameters of methyl groups $S^2_{axis}$ in (A) 1918, (B) PR8, (C) Ud, and (D) VN NS1s.

Supplementary Figure 15

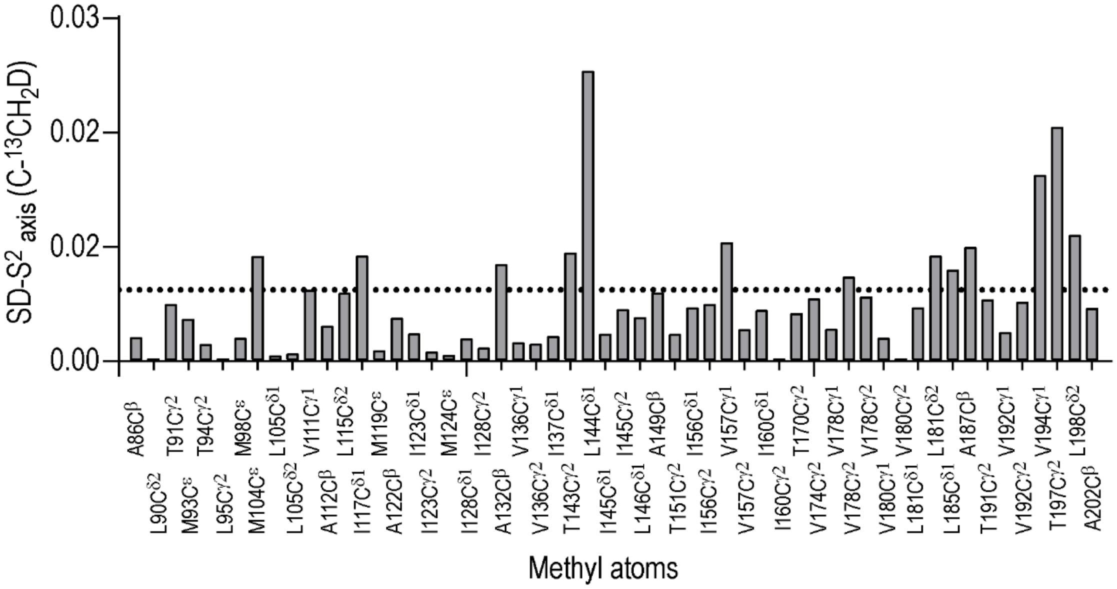

**Supplementary Figure 15.** Standard deviation of $S^2_{axis}$ values of NS1 proteins. Mutated residues were excluded from the calculation. Dotted line represents Q3 of standard deviations.
